# Quantifying between-species differences in morphological integration patterns using two-block PLS: an R implementation and sensitivity analysis

**DOI:** 10.64898/2025.12.04.692052

**Authors:** Mikel Arlegi

## Abstract

Morphological integration is commonly quantified within species using covariance or correlation structures, and two-block partial least squares (PLS) has become a standard tool for assessing covariation between anatomical modules. However, most applications focus on a global PLS axis for a pooled sample, and there is no simple, widely used procedure to compare whether integration patterns differ between species in the same PLS space. Here I present an angle-based method and an accompanying R function, Integration_Patterns, that quantify between-species differences in integration patterns based on the orientation of the first eigenvector of the within-species covariance matrix in PLS1–PLS1 space. For each species, the method extracts PLS1 scores for two modules, computes the covariance matrix, retrieves the principal eigenvector representing the main axis of covariation, and then measures the angle (0–90°) between these axes across species. Small angles indicate nearly parallel integration patterns, whereas larger angles indicate divergent or orthogonal patterns.

To evaluate the behaviour and robustness of the estimator, I conducted a simulation-based sensitivity analysis using species-specific PLS1 scores. Pairs of species were simulated under known angular separations (0°, 30°, 45°, 60°, 90°) and a range of per-species sample sizes (*n* = 3 to 200). For each combination, 100 replicate datasets were generated and the between-species angle was estimated. The results show that the estimator is biased and highly variable for very small samples (*n* ≤ 10), but converges rapidly to the true angle with increasing sample size. For *n* ≈ 20–30 individuals per species, the mean estimated angle is close to the true value and the standard deviation is substantially reduced across all scenarios, including orthogonal patterns (90°). These findings provide practical guidance on sample-size requirements when comparing integration patterns across species.

## 1 Introduction

Morphological integration refers to the tendency of traits to co-vary due to developmental, functional or genetic factors (Olson & Miller, 1958). Analyses of integration have become central to evolutionary morphology because they reflect how variation is structured, how modules respond to selection, and how constraints and opportunities shape morphological evolution (Cheverud, 1996; Mitteroecker & Bookstein, 2007; Mitteroecker et al., 2012; Klingenberg, 2014; Adams & Collyer, 2016). Consequently, integration has played an important role in understanding how suites of traits evolve across phylogenies and ecological contexts (e.g., Rolian, 2009; Grabowski et al., 2011; Gómez-Robles & Polly, 2012; Goswami et al., 2014). In geometric morphometrics, integration is often assessed using covariance matrices of Procrustes-aligned coordinates, PLS between anatomical modules, and measures of covariation strength such as r-PLS (Klingenberg, 2008; Goswami & Polly, 2010; Adams & Collyer, 2016: Arlegi et al., 2018, 2022).

Two-block PLS has been particularly influential for studying integration between pairs of modules. In this context, PLS identifies linear combinations of variables in each block that maximise covariation between them (Rohlf & Corti 2000). The first PLS axis (PLS1) captures the strongest mode of covariation, and its associated scores describe how specimens vary jointly across both modules. Most empirical studies interpret the strength of integration via r-PLS and visualise PLS1 shape changes, often pooling specimens across species or populations.

However, many evolutionary questions require comparing how integration patterns differ among species, not just whether there is significant integration in a pooled sample. One may wish to know whether two species share a similar cranium–cervical integration pattern, whether integration has been reoriented along a different axis in a particular clade, or whether differences in integration are associated with changes in posture, locomotion or ecological niche. Current workflows offer limited tools for such between-species comparisons. Although separate PLS analyses can be fitted for each group or group-specific covariance matrices can be compared indirectly, there is no simple or standardised procedure for quantifying whether integration patterns are parallel or divergent across species within a shared PLS space (Mitteroecker & Bookstein 2007; Klingenberg 2009, 2013; Goswami & Polly 2010; Zelditch et al. 2012; Adams & Collyer 2016).

Here I address this limitation by introducing an approach that extracts, for each species, the dominant axis of covariation between modules and quantifies how these axes differ across species using angular distances and permutation-based significance tests. The method focuses on the geometry of integration patterns in the bivariate PLS1–PLS1 score space, where each species exhibits its own within-species covariance structure. This structure can be summarised by the first eigenvector of the covariance matrix, which represents the species-specific axis of maximal covariation. Comparing the orientation of these axes provides a direct measure of between-species differences in integration patterns: small angles indicate nearly parallel covariation patterns, whereas larger angles reflect increasingly divergent, up to orthogonal, patterns. Conceptually, this is analogous to comparing allometric trajectories between groups, where parallel trajectories imply similar underlying responses and divergent trajectories suggest distinct patterns of covariation.

In this work, I formalise this approach by presenting an angle-based framework for comparing species-specific integration patterns and an accompanying R implementation, Integration_Patterns. The method is embedded within the two-block PLS framework of R package Morpho v.2.13 (Schlager, 2017), and provides a practical pipeline for extracting species-specific axes of covariation and quantifying their divergence. I evaluate its statistical behaviour through a simulation-based sensitivity analysis across a range of true angular differences and sample sizes, and illustrate its application using a controlled example with five species exhibiting known integration axes.

## 2 Methods

### 2.1 Statistical framework

The method assumes that a two-block PLS analysis has been carried out on two sets of shape variables (e.g., Procrustes coordinates or other multivariate descriptors) representing two anatomical modules. A single PLS is fitted to the full dataset so that all individuals, and thus all species, are embedded in a common PLS space. Two-block PLS is performed using the function pls2B from the R package Morpho v.2.13 (Schlager, 2017), which decomposes the covariance between the X- and Y-blocks into pairs of latent variables (PLS axes). The first axis (PLS1) captures the maximal shared covariation between the two modules, and for each individual, pls2B returns scores for the X-block (XScores) and the Y-block (YScores). In what follows, these PLS1 scores are treated jointly as a bivariate point in PLS1–PLS1 space.

The function Integration_Patterns assumes that the user provides two arrays or matrices X and Y containing matching individuals, and a factor or character vector species of length equal to the number of individuals, indicating species identity. Individuals are pooled across species for the PLS, ensuring a shared latent space, and species-specific patterns are then characterised within this common space. The method requires at least three individuals per species to compute within-species covariance matrices, although larger samples are needed for stable estimation, as explored in the sensitivity analysis.

For each species, the paired PLS1 scores from the two modules are treated as a bivariate dataset describing how individuals vary jointly in the PLS1–PLS1 space. The covariance matrix of the PLS1 scores summarises the within-species pattern of covariation, and its first eigenvector defines the dominant axis of joint variation. Because eigenvectors represent axes rather than oriented vectors, their sign is arbitrary and they are treated as undirected lines.

To compare integration patterns between species s_1_ and s_2_, with principal eigenvectors v_1_,s_1_ and v_1_,s_2_, the raw angle between them is

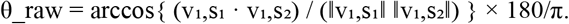

Since opposite orientations correspond to the same axis, θ_raw and 180° − θ_raw represent equivalent separations. The between-species angle is therefore

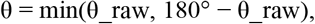

which constrains θ to the interval [0°, 90°]. Small values indicate nearly parallel integration patterns, whereas values approaching 90° reflect strongly divergent or orthogonal patterns. Pairwise angles for all species are assembled into an S × S matrix summarising between-species differences in integration patterns.

### 2.2 Implementation in R

The R function Integration_Patterns automates this workflow. Given X, Y and species, it (i) runs Morpho::pls2B to obtain PLS1 scores for both blocks, (ii) constructs a bivariate matrix of PLS1 scores and splits it by species, (iii) for each species with sufficient sample size, computes the covariance matrix, eigenvalues and eigenvectors, retaining the first eigenvector, and (iv) computes all pairwise angles between species-specific eigenvectors. The function returns a list containing the PLS result, a data frame of PLS1 scores with species labels, species-wise covariance matrices and eigenvectors, and the angle matrix.

For any pair of species, a permutation procedure can be used to evaluate whether the observed angular difference is greater than expected under a null hypothesis of no species-specific integration pattern. The null model assumes that individuals are exchangeable with respect to species, given their PLS1 scores. For a given pair of species, the PLS1 scores are pooled, species labels are randomly permuted while keeping the scores fixed, and the angle between the principal axes of the two permuted groups is recomputed. Repeating this procedure n_perm times yields a null distribution of angles under the hypothesis that both species share the same integration axis in PLS1–PLS1 space. Large observed angles relative to the null distribution indicate that the two species exhibit significantly different integration patterns, whereas small angles may not differ from what is expected by chance given the within-species variation.

### 2.3 Simulation-based sensitivity analysis

To evaluate the statistical properties of the angle estimator, I conducted a simulation study directly in the PLS1–PLS1 space, where the method operates. Instead of simulating landmark configurations and re-running PLS for each replicate, the simulations generated bivariate PLS1 score distributions that reproduce the covariance structure expected for two anatomical modules exhibiting a given integration pattern. Each species’ scores were drawn from a bivariate normal distribution whose covariance ellipse was oriented at a specified angle, thereby imposing a known integration axis for that species.

In each scenario, two species were simulated: a reference species with its integration axis fixed at 0°, and a comparison species whose axis was set to one of five values (0°, 30°, 45°, 60° and 90°), representing true between-species differences ranging from none to complete orthogonality. Sample size per species varied from *n* = 3 to *n* = 200 to assess how the estimator behaves under conditions of limited or abundant replication. For every combination of true angle and sample size, 100 replicate datasets were generated. In each replicate, individuals were simulated for both species under their assigned covariance structures. Species-specific covariance matrices were then estimated from the paired PLS1 scores, the principal integration axis was extracted for each species, and the between-species angle was computed following the axial definition described earlier. For each scenario, the mean estimated angle and its standard deviation across replicates were summarised. These values provide a clear indication of how rapidly the estimator converges on the true angle and how its variability declines as sample size increases.

### 2.4 Example with five species simulated data

To illustrate how Integration_Patterns can be applied to multivariate data, I generated a simple simulated dataset comprising five hypothetical species (A–E), each with *n* = 100 individuals. The goal of this example was not to emulate the full complexity of 3D landmark data, but to produce interpretable PLS1 score distributions whose integration patterns differ in known ways among species.

For each species, paired data representing two anatomical modules were simulated as bivariate PLS1 scores drawn from multivariate normal distributions with strong covariance and species-specific orientation of the major axis. The true covariation angles were set to 30° (A), 45° (B), 60° (C), 75° (D) and 90° (E). In the PLS1–PLS1 plane, each species thus formed an elongated elliptical cloud oriented at its assigned angle. Species with similar angles (A vs B, C vs D) generated similar patterns, whereas species with more divergent orientations (e.g., A vs E) showed clearly different integration axes.

In this example, I worked directly in PLS1–PLS1 space. The Integration_Patterns workflow was applied as it would be after a real PLS: individuals were assigned to species, bivariate PLS1 scores were split by species, species-specific covariance matrices were computed, principal eigenvectors were extracted, and all pairwise angles were calculated using the acute (0–90°) definition. For selected pairs of species, permutation tests were used to assess the significance of the observed angle by randomising species labels and recomputing the angle, yielding *p*-values based on the null distribution of angles under no species-specific integration pattern.

## 3 Results

### 3.1 Sensitivity analysis

Results are represented in Figure 1 and Table S1. When both species shared the same underlying integration pattern (Δθ = 0°), the estimated angle between species was non-zero for small sample sizes due to sampling variation in the estimated covariance matrices and eigenvectors. At *n* = 3 individuals per species, mean estimated angles were large and highly variable. At *n* = 5 and *n* = 10, mean angles decreased but remained clearly above 0°, reflecting instability in the estimation of principal axes at low sample sizes. As sample size increased, both bias and variance declined. At *n* ≈ 20, the mean estimated angle was around 9° with moderate standard deviation. By *n* ≈ 30, mean angles were approximately 7° with reduced variance, and at *n* = 50 they were close to 5° with relatively narrow dispersion. For *n* ≥ 100 individuals per species, mean angles fell below 4°, and at *n* = 200 the estimates were very close to 0° with tight variability (Fig. 1). These results show that, even when integration patterns are truly identical, finite sample sizes yield spurious between-species angles, but the estimator converges towards 0° as n increases, and for *n* on the order of 30 or more the residual between-species angle under a parallel pattern is modest and precise.

**Figure 1.**
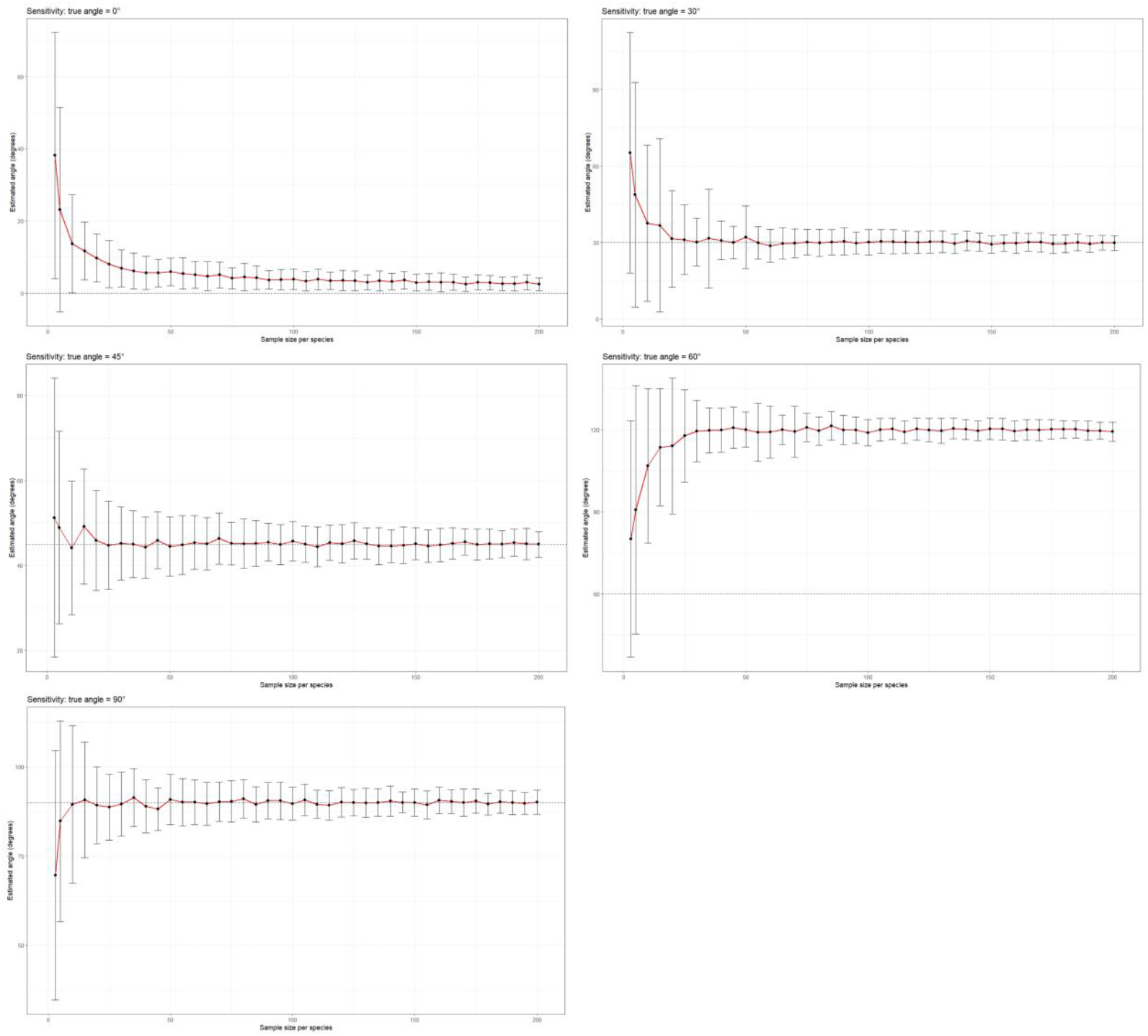
Sensitivity of the angle estimator across a range of true between-species differences and sample sizes. Sensitivity curves for five simulation scenarios, where the true angular separation between species-specific integration axes was 0°, 30°, 45°, 60° and 90°. For each sample size (3–200 individuals per species), 100 replicate datasets were simulated, and the mean estimated angle (red line) and its standard deviation (vertical error bars) were computed. The dashed horizontal line indicates the true angle in each scenario.

Under moderate differences in the true integration axis (Δθ = 30°, 45° and 60°), the estimator converged rapidly to the correct value as n increased. For Δθ = 30°, mean estimated angles were already close to the true value by *n* = 10, albeit with substantial variance, and stabilised around 30° with decreasing variance as *n* increased. For Δθ = 45°, a similar pattern was observed: estimates were centred near 45° even at relatively small *n*, and the spread around the true value declined steadily with increasing sample size. For Δθ = 60°, mean estimates were close to 60° for *n* ≥ 10 and remained centred on that value while the standard deviation decreased as *n* grew (Fig. 1). Across these moderate-angle scenarios, the estimator was effectively unbiased for *n* ≥ 20–30 and exhibited steadily decreasing variance with increasing sample size. Differences of 30–60° are therefore reliably estimated with sample sizes that are realistic for many comparative morphological studies, provided that each species contributes at least 20–30 individuals.

The scenario with Δθ = 90° represents the maximal divergence in PLS1–PLS1 space. For very small sample sizes, the estimator was again noisy, with mean angles substantially below 90° and high variance at *n* = 3 and *n* = 5. At *n* = 10, the mean estimated angle was much closer to the true value but still showed considerable spread across replicates. With increasing sample size, mean angles converged towards 90° and variability decreased. Around *n* = 20–30, estimates were consistently centred near 90° with moderate variance, and for *n* ≥ 100 the distribution of estimates tightened substantially. These results indicate that, even in the extreme case of orthogonal integration patterns, the estimator converges reliably to the true angle and becomes precise once sample sizes reach *n* ≈ 20–30.

Taken together, the simulations yield three general conclusions. First, the angle estimator is unstable and biased at very small sample sizes (*n* ≤ 10), both under parallel and divergent patterns, because within-species covariance matrices are poorly estimated and principal axes may appear spuriously rotated. Second, for moderate to large sample sizes (*n* ≈ 20–30 and above), the estimator is approximately unbiased and its variance declines markedly. For true angles of 30°, 45° and 60°, mean estimates are very close to the true values, with standard deviations typically below 10° at *n* ≈ 30 and around 3–4° at *n* = 200. For the parallel case, spurious between-species angles shrink to only a few degrees as *n* increases. Third, even in the orthogonal scenario (90°), the estimator is well behaved, converging towards 90° and achieving high precision for *n* ≥ 30.

### 3.2 Example

The results of this analysis are represented in Table 1. The five-species example further illustrates the performance of the method when integration axes are known a priori. Species A–E were assigned true integration angles of 30°, 45°, 60°, 75° and 90°, respectively, so that expected pairwise differences ranged from small (A–B: 15°) to large (A–E: 60°). The estimated angles closely matched these theoretical differences. Species with similar true orientations (A vs B; C vs D) showed small angular distances (A–B ≈ 15.1°, C–D ≈ 9.0°). Intermediate contrasts (e.g., B–C, 45° vs 60°) yielded moderate angles (≈ 22.7°). Contrasts involving strongly divergent species (A–E, 30° vs 90°; B–E, 45° vs 90°) produced large angles of ≈ 61.0° and ≈ 46.0°, respectively. These patterns indicate that the method reliably recovers both the ordering and magnitude of between-species differences in integration axes.

**Table 1.** Pairwise angular differences and permutation-based significance tests for the simulated five-species example. Upper matrix, angles (°) between species-specific integration axes in PLS1–PLS1 space. Lower matrix: *p*-values from permutation tests assessing whether observed angles exceed those expected under random species assignments.

|  | **A** | **B** | **C** | **D** | **E** |
| --- | --- | --- | --- | --- | --- |
| **A** |  | 15.081 | 37.767 | 46.740 | 61.036 |
| **B** | 15.081 |  | 22.686 | 31.659 | 45.955 |
| **C** | 37.767 | 22.686 |  | 8.973 | 23.269 |
| **D** | 46.740 | 31.659 | 8.973 |  | 14.296 |
| **E** | 61.036 | 45.955 | 23.269 | 14.296 |  |
| **A** |  | 0.003 | 0.000 | 0.000 | 0.000 |
| **B** | 0.003 |  | 0.000 | 0.000 | 0.000 |
| **C** | 0.000 | 0.000 |  | 0.017 | 0.000 |
| **D** | 0.000 | 0.000 | 0.017 |  | 0.000 |
| **E** | 0.000 | 0.000 | 0.000 | 0.000 |  |

Permutation tests applied to the five-species example further confirm that the method detects genuine differences in integration patterns. Under the null model in which species labels are randomly reassigned, small observed angles between species with similar true axes (A–B, C–D) produced relatively higher *p*-values (e.g., C–D: *p* = 0.017), consistent with the possibility that such small differences could occasionally arise by chance. In contrast, large observed angles (A–C, A–D, A–E, B–D, B–E, C–E) yielded extremely low *p*-values (*p* ≈ 0 or < 0.003), indicating that such strong divergences almost never occur under random assignment. All tests involving species E, whose true axis differed most strongly from the others, showed highly significant differences (*p* ≈ 0).

The overall pattern of *p*-values matches the simulation design: species with similar integration patterns are statistically indistinguishable, whereas species with divergent covariation axes are identified as significantly different. This example demonstrates that the method accurately reconstructs species-specific integration axes from covariance matrices of PLS scores, that pairwise angular distances correspond closely to true underlying differences, and that permutation tests correctly differentiate between small, moderate and large differences without inflating false positives. The method is sensitive enough that even modest differences (e.g., 15° between A and B) are recovered, and large differences (≥ 45°) are detected with strong support.

## Discussion

This preprint introduces an angle-based framework and an R implementation for quantifying between-species differences in morphological integration patterns within a two-block PLS context. Rather than focusing solely on the global strength of integration using two-block PLS, the method characterises how species differ in the orientation of their main covariation axis in PLS1–PLS1 space. The main idea is to summarise within-species covariation by the first eigenvector of the PLS1–PLS1 covariance matrix and to compare these vectors between species through their angular separation.

The method is simple, interpretable and compatible with existing geometric morphometric workflows. It builds on established tools such as Morpho’s pls2B (Schlager, 2017) by adding an explicit layer for between-species comparison of integration patterns. Conceptually, this is analogous to comparing allometric trajectories across groups, but applied to the main axis of covariation between two modules rather than to size–shape relationships. Because eigenvectors represent axes without inherent direction, angles are treated as axial and folded into the 0–90° range, ensuring that parallel but oppositely oriented vectors are recognised as representing the same integration pattern.

The between-species angle has a straightforward biological interpretation. Values near 0° indicate that species share a nearly parallel integration pattern: the main axis of covariation is aligned and differences may reflect scale or variance magnitude more than orientation. As the angle increases, integration patterns are interpreted as increasingly divergent. At 90°, the principal axes are orthogonal, indicating maximal difference in how the two modules co-vary within each species. In comparative contexts, such angles can be used to identify clusters of species with similar integration patterns, test hypotheses about shifts in integration associated with functional or ecological transitions, and assess whether pooling species or analysing residuals is appropriate, particularly when evaluating whether size or another covariate should be removed.

The five-species example demonstrates that the method behaves as expected when the true integration axes are known. In that example, species A–E were assigned covariation angles of 30°, 45°, 60°, 75° and 90° in PLS1–PLS1 space. The estimated pairwise angles closely mirrored these theoretical differences: species with similar true orientations (A vs B; C vs D) showed small angular distances, while contrasts involving strongly divergent species (e.g., A vs E) yielded large angles close to the expected 60°. Permutation tests confirmed that these large angular differences are highly unlikely under a null model of no species-specific integration pattern, whereas smaller angles between species with similar true axes were less extreme and, in some cases, not strongly significant. Together, these results indicate that the method can recover both the rank order and the magnitude of between-species differences in integration patterns and that the permutation framework provides a coherent way to assess their statistical support.

The sensitivity analysis provides practical guidance on the sample sizes required for reliable estimation. Under a true parallel pattern (0°), small sample sizes yielded non-zero estimated angles due to sampling variation in covariance matrices and eigenvectors. This highlights a generic limitation of covariance-based approaches: with *n* ≤ 10 individuals per species, covariance summaries are inherently unstable and apparent between-species differences in integration patterns may be dominated by noise. However, the simulations also show that the estimator is well behaved for larger samples. For moderate angular differences (30°, 45°, 60°), estimated angles were approximately unbiased from *n* ≈ 20–30 onwards, with variance declining rapidly as n increased. For extremely divergent patterns (90°), convergence to the true value was also rapid, although variance remained higher at very small *n*.

A practical implication is that studies aiming to compare integration patterns between species should, where possible, aim for at least 20–30 individuals per species. Below this threshold, it is still possible to compute angles, but they should be interpreted with caution and ideally accompanied by uncertainty estimates or sensitivity checks (such as subsampling or simulations tailored to the empirical design). The five-species example reinforces this point, the method can clearly distinguish large differences in integration axes when sample sizes are adequate, but similar patterns could be much harder to detect robustly with very small samples.

The method presented here complements existing approaches to morphological integration. Two-block PLS remains the primary tool for assessing covariation between modules (Rohlf & Corti, 2000), and r-PLS derived from pooled or group-specific analyses remains useful for quantifying the strength of integration (Adams & Collyer, 2016). However, *r*PLS alone does not capture differences in the orientation of integration patterns among species, two taxa may show similar levels of integration strength but differ in how variation is structured. By focusing on eigenvector orientation, Integration_Patterns adds a new dimension to integration analysis, making it possible to formally assess whether integration patterns are effectively parallel, moderately divergent or strongly distinct across taxa.

Several limitations and potential extensions deserve mention. First, the current implementation focuses on the first eigenvector of the within-species covariance matrix in the PLS1–PLS1 plane. This captures the dominant integration pattern but ignores secondary axes that may be biologically relevant in some cases. Extending the approach to compare higher-dimensional subspaces or multiple PLS axes would be a natural next step, albeit at the cost of more complex interpretation. Second, the present implementation treats species as independent units and does not incorporate phylogenetic information. In an evolutionary framework, it would be desirable to map integration pattern shifts onto a phylogeny, reconstruct ancestral integration axes and test whether changes in integration are associated with particular branches, nodes or adaptive transitions. Third, the method assumes that the PLS1–PLS1 space is an appropriate summary of the covariation structure. In many empirical datasets, PLS1 captures a large proportion of the covariation between modules, but this may not always hold, and diagnostics of the dimensionality and structure of covariation remain important.

Overall, this work introduces a simple, interpretable framework for comparing morphological integration patterns between species in a two-block PLS context. By summarising within-species covariation in PLS1–PLS1 space and quantifying between-species differences as angles between principal eigenvectors, the method offers a direct way to assess whether integration patterns are parallel or divergent across taxa. Simulation-based sensitivity analyses show that the estimator is biased and variable for very small sample sizes but converges rapidly to the true angle with *n* ≈ 20–30 individuals per species and above, and the five-species example confirms that the method can recover known differences in integration axes and detect them as statistically significant where appropriate. Integration_Patterns is intended as a general tool for evolutionary morphologists interested in how integration evolves, how it differs among species, and how it relates to functional, ecological or developmental factors.

## Acknowledgements

I gratefully acknowledge the support of the British Academy through the British Academy International Fellowship (grant IF23/100647). I also acknowledge support from the European Union’s Horizon Europe research and innovation programme under the Marie Skłodowska-Curie Postdoctoral Fellowship CR-EVOL (grant agreement EU-MSCA-101108040).

## Data availability

The code for the Integration_Patterns function is available on GitHub (IntegrationPatterns/README.md at main · Mikel971/IntegrationPatterns · GitHub) and archived on Zenodo (DOI: 10.5281/zenodo.17623868).

## Supplementary Information

Table S1 Numeric results obtained for the sensitivity analysis represented in Figure 1.

